# Unilateral movement preparation causes task-specific modulation of TMS responses in the passive, opposite limb

**DOI:** 10.1101/304410

**Authors:** Lilian Chye, Stephan Riek, Aymar de Rugy, Richard G. Carson, Timothy J. Carroll

## Abstract

Corticospinal excitability is modulated for muscles on both sides of the body during unilateral movement preparation. For the effector, there is a progressive increase in excitability, and a shift in direction of muscle twitches evoked by transcranial magnetic stimulation (TMS) toward the impending movement. By contrast, the directional characteristics of excitability changes in the opposite (passive) limb have not been fully characterized. Here we assessed how preparation of voluntary forces towards four spatially distinct visual targets with the left wrist alters muscle twitches and motor evoked potentials (MEPs) elicited by TMS of left motor cortex. MEPs were facilitated significantly more in muscles homologous to agonist rather than antagonist muscles in the active limb, from 120 ms prior to voluntary EMG onset. Thus, unilateral motor preparation has a directionally-specific influence on pathways projecting to the opposite limb that corresponds to the active muscles rather than the direction of movement in space. The directions of TMS-evoked twitches also deviated toward the impending force direction of the active limb, according to muscle-based coordinates, following the onset of voluntary EMG. The data indicate that preparation of a unilateral movement increases task-dependent excitability in ipsilateral motor cortex, or its downstream projections, that reflect the forces applied by the active limb in an intrinsic (body-centered), rather than an extrinsic (world-centered), coordinate system. The results suggest that ipsilateral motor cortical activity prior to unilateral action reflects the state of the active limb, rather than subliminal motor planning for the passive limb.

## Key point summary

- Activity in the primary motor cortices of both hemispheres increases during unilateral movement preparation, but the functional role of ipsilateral motor cortex activity is unknown.
- Ipsilateral motor cortical activity could represent subliminal “motor planning” for the passive limb. Alternatively, it could represent the state of the active limb, to support coordination between the limbs should a bimanual movement be required.
- Here we assessed how preparation of forces toward different directions, with the left wrist, alters evoked responses to transcranial magnetic stimulation of left motor cortex.
- Preparation of a unilateral movement caused excitability increases in ipsilateral motor cortex that reflected forces produced with the active limb in an intrinsic (body-centered), rather than an extrinsic (world-centered), coordinate system.
- These results suggest that ipsilateral motor cortical activity prior to unilateral action reflects the state of the active limb, rather than subliminal motor planning for the passive limb.

## Introduction

Executing even an apparently simple movement, such as picking up an object, requires non-trivial sensorimotor integration. The brain must derive information from multiple sensory systems that is defined in various frames of reference; including eye-based (visual), head-based (vestibular), joint-based (skin and joint afferent) and muscle-based (muscle afferent) receptor systems (Andersen *et al*., 1993; Sabes, 2011). This sensory information must be integrated to provide estimates of the state of the body and relevant features of the environment, and must ultimately be transformed into a set of motor commands (Andersen *et al*., 1993; Buneo & Andersen, 2006; Sabes, 2011). During motor preparation, the firing rates of neurons in multiple parietal and frontal regions that contribute to sensorimotor control correlate with directionally specific features of forthcoming movement (Buneo & Andersen, 2006). Many of these areas appear to represent the direction of impending movement in more than one coordinate system (Sabes, 2011). For example, the activity of separate sub-populations of cells in primary motor cortex (M1) may correspond to either the specific muscles that will be activated, or to the extrinsic direction of the motion that will be generated irrespective of the muscles involved (Kakei *et al*., 1999). Indeed, even among corticomotoneuronal cells in M1 which project monosynaptically to motoneurons in the spinal cord, the directions of movement associated with the highest firing rates may be distinct from the pulling directions of the muscles that they actuate (Griffin *et al*., 2015). Thus, it appears that neural activity in M1 reflects both intrinsic and extrinsic movement features (Kalaska & Crammond, 1992; Scott *et al*., 2001; Sabes, 2011).

During voluntary unilateral movements of the upper limb, the M1 contralateral to the active limb (subsequently referred to as M1_contra_) plays an instrumental role via the ~80% corticospinal fibres that cross to the contralateral hemicord at the pyramidal decussation (Kertesz & Geschwind, 1971; Siegel & Sapru, 2011; Levy, 2013). However, there is increasing evidence that the excitability of circuits in M1 ipsilateral to the active limb (subsequently referred to as M1_ipsi_) is also modulated during unilateral movement (Kim *et al*., 1993; Duque *et al*., 2005; McMillan *et al*., 2006; Perez & Cohen, 2008; Duque & Ivry, 2009; Duque *et al*., 2010; Hinder *et al*., 2010; Lee *et al*., 2010; Howatson *et al*., 2011; Verstynen & Ivry, 2011; Carson & Ruddy, 2012; Bütefisch *et al*., 2014; Chiou *et al*., 2014; Perez *et al*., 2014). Neural activity in M1_ipsi_ of non-human primates also correlates with movement parameters such as endpoint direction and velocity during unilateral movement preparation and execution (Ganguly *et al*., 2009), to the extent that unilateral limb movements can even be “decoded” from M1_ipsi_ firing rates (Ganguly & Poo, 2013). However, the functional purpose of this ipsilateral activity in M1 is still unclear: does it represent subliminal “motor planning” for the passive limb, or does it represent information about the current or future state of the active limb (e.g. via efference copy) to support the capacity for coordination between the limbs should a bilateral movement be required?

One way to address this question is to consider the coordinate frames to which activity in M1_ipsi_ corresponds. As the body is mirror symmetric, when registered in an extrinsic coordinate system, activation of homologous muscles that generate motion about the frontal axes moves the limbs in opposite directions (e.g. leftward or rightward in extrinsic space). Conversely, the direction of movement brought about by the activation of homologous muscles is always the same for both limbs according to an intrinsic coordinate system. Thus, if activity in the ipsilateral M1 corresponds to subliminal motor planning for the passive limb, it should predominantly reflect the task goal in extrinsic coordinates (i.e. resulting in increased excitability for muscles that would move the passive limb to the goal). Conversely, ipsilateral M1 activity that corresponds to the preparatory state of the active limb should at least partly reflect the task goal according to intrinsic coordinates (i.e. resulting in increased excitability for muscles homologous to those due to be recruited in the active limb).

Previous work indicates that the excitability of projections to individual muscles in the passive limb is modulated during preparation of movements performed by the opposite limb (Leocani *et al*., 2000; Duque *et al*., 2005; McMillan *et al*., 2006; Duque *et al*., 2008; Duque & Ivry, 2009; Duque *et al*., 2010). Leocani et al. (2000) reported that the excitability of projections to homologous muscles was reduced during unimanual preparation, when activation of these muscles would have produced motion toward the body midline. On the basis of a series of experiments in which the posture of the arm was manipulated, Duque et al. (2005; 2008; 2010) further concluded that the inhibition of projections from M1_ipsi_ is associated with the direction of motion with respect to the body midline (toward or away), rather than the specific muscles involved in moving the effector in a particular direction. In these studies however, the required direction of motion was specified by symbolic cues rather than spatial targets. This situation might favor body-referenced over extrinsic neural representations, since there is no obligation for any motor preparatory activity to be represented according to an extrinsic reference frame in the absence of an extrinsic spatial target. Conversely, McMillan et al. (2006) reported increases in the excitability of projections to muscles (in the opposite limb) that had lines of action consistent with the extrinsic direction of motion required to acquire a pre-cued spatial target. This observation suggests that ipsilateral M1 activity during the preparation of visually-guided unilateral actions may also reflect the extrinsic direction of the forthcoming movement in some circumstances. In each of these studies however, there were interleaved trials in which the passive limb was required to move, either in isolation or as part of a bimanual action. The task context was therefore one in which movements of each limb were either to be selected or inhibited at short latency.

The variety of task contexts and methodology used in previous work permit a number of plausible hypotheses about the fundamental nature of activity in M1_ipsi_. One possible framework for interpreting the different results, i.e. that ipsilateral activity can have either intrinsic or extrinsic representations depending on context, is that ipsilateral activity represents the extrinsic goal of the active limb when there is a definite spatial target, but that in the absence of a spatial target, there is a default tendency for activity to mirror the active limb according to a frame of reference defined by the body midline. However this conjecture fails to accommodate the fact that in previous work, both limbs were required to prepare for movement in some trials. In the current study, we therefore set out to examine responses to TMS of M1_ipsi_ in a strictly unilateral movement context that involved an extrinsic spatial target. This situation resembles natural conditions in which people make unilateral movements to interact with physical objects, since the muscle activity generated to interact with an object has a well-defined extrinsic spatial goal. Our aim was to distinguish whether changes in the excitability of corticospinal projections to the passive limb reflect the intrinsically– or extrinsically-defined direction of force produced by the effector limb under these conditions. More specifically, if preparation of a strictly unilateral movement to a spatial target increases the size of responses following stimulation of ipsilateral motor cortex according to in an intrinsic coordinate system, it would support the hypothesis that ipsilateral motor cortical activity prior to unilateral action reflects the state of pathways projecting to the active limb, rather than subliminal motor planning for the passive limb.

## Methods

Twelve right-handed participants (11 males and 1 female; aged between 20 and 37 years old) with no recent wrist, elbow or shoulder injuries volunteered for the study. Right-handedness was confirmed with Edinburgh Handedness Inventory (Oldfield, 1971). A medical questionnaire was used to screen the participants for neurological disorders and contraindications in relation to the application of TMS (Rossi *et al*., 2011; Groppa *et al*., 2012). The study was approved by the Medical Research Ethics Committee of The University of Queensland (HMS13/0506). All participants were briefed on the experimental procedures and gave written informed consent prior to the experiment, which conformed to the Declaration of Helsinki.

### Experimental protocol

The participants were required to attend a single experimental session in which they performed a choice reaction time, isometric, force aiming task with the left wrist. The aiming task required them to move a cursor that represented the resultant force exerted at the wrist joint in 2 degrees of freedom (ab-adduction versus flexion-extension) to one of four targets, which appeared in random order along the cardinal axes (i.e. 0°, 90°, 180°, 270°; Figure 1B). Although the task involves only very minor displacement of the limb end-point, it does require shortening of muscle fibres (and concomitant lengthening of tendons), and motion of the cursor that represents force magnitude. Thus, for simplicity of expression, we refer to the isometric actions produced in this task as “movements” throughout the paper. We simultaneously measured force twitches evoked by TMS at the passive right wrist in some trials (see below for details). Prior to the beginning of the experiment, participants completed two blocks (48 trials) of the choice reaction time task as familiarisation. Each individual’s average reaction time was estimated from the familiarisation trials to define the timing of TMS for the main experiment.

**Figure 1.**
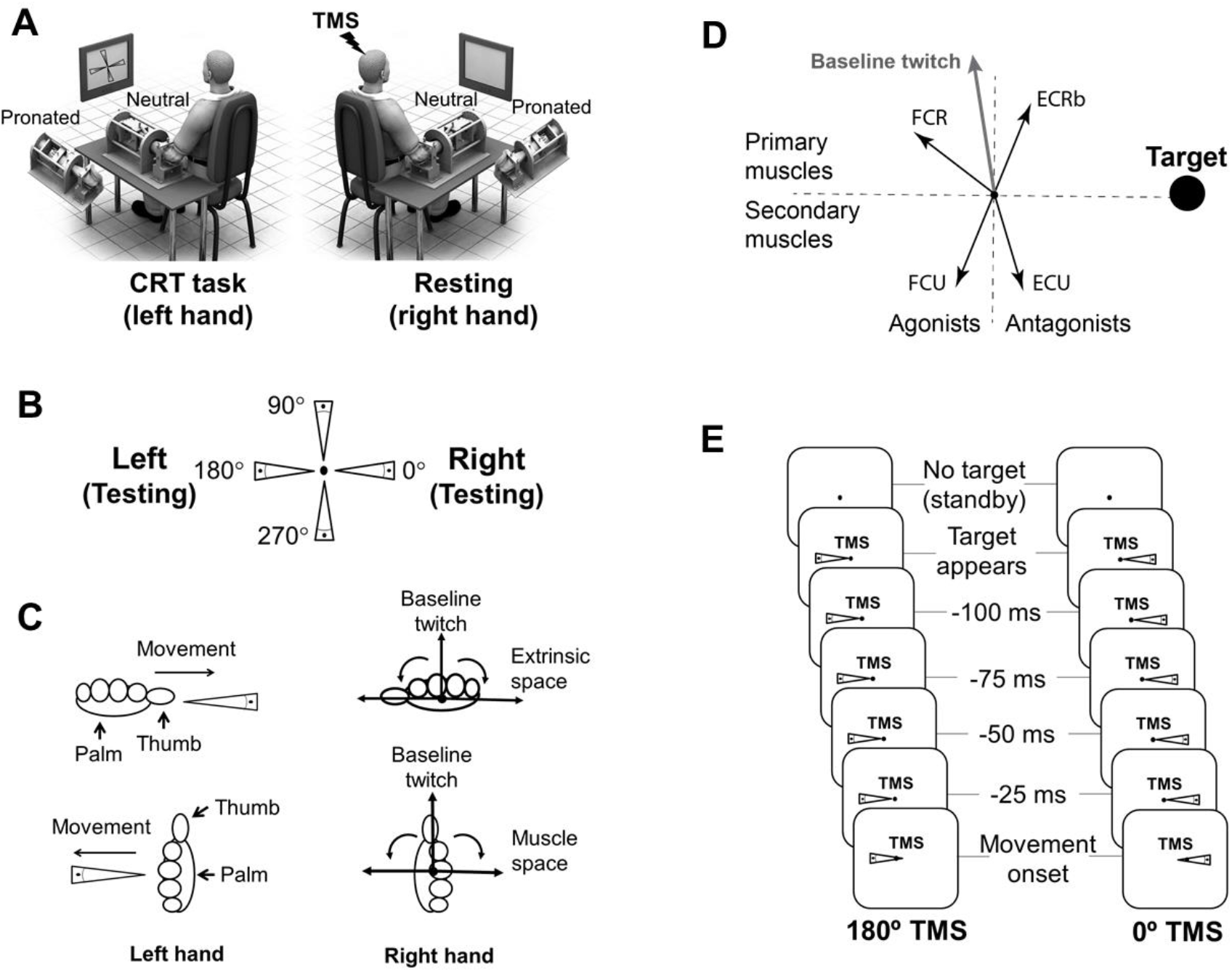
**(A)** Left figure shows participants making movements with their left wrist toward one of four targets in the choice reaction time task. Right figure shows that TMS was applied the left motor cortex to evoke twitches from the right wrist at rest, during movement preparation and upon movement onset of the left wrist. **(B)** Illustration of the four-alternative choice reaction time task. The targets appeared in a randomised order and participants made quick isometric forces upon target appearance with their left hand. **(C)** Schematic representation of the TMS-evoked twitch directions from resting right wrist, during preparation of a movement with the left wrist, toward horizontal targets in pronated and neutral hand positions. The figures on the left illustrate pronated and neutral hand positions with movement direction toward a horizontal target, i.e. 0° and 180° targets respectively. The right figures illustrate how the twitch direction changes in the passive limb would correspond to active force in the opposite limb according to different reference frames. Note that only the outer reference frame is labelled for clarity; if an outward direction corresponds to an extrinsic reference frame, then an inward direction necessarily corresponds to a muscle-based reference frame. Note also that a muscle space frame of reference is indistinguishable from a body midline-based reference frame in this design. **(D)** Schematic representation of the way muscles were grouped for analysis with the forearms in the neutral orientation. The plot shows the pulling directions of muscles in the passive right wrist, taken from (de Rugy *et al*., 2012), when the active left limb exerted force toward the 0° target (horizontal right target). The figure illustrates that the FCR and FCU muscles are homologous to the agonists in the active limb, and the ECR and ECU are homologous to the antagonists. The muscles were further categorised as “primary” or “secondary” on the basis of whether their pulling directions aligned with the baseline twitch direction. Primary agonists are defined as muscles with pulling directions lying between the target direction and the baseline twitch direction. Secondary agonists are defined as muscles with pulling directions aligned with the target (i.e. within 90 degrees), but away from the baseline twitch direction (more than 90 degrees away). Antagonist muscles are similarly specified as primary or secondary. **(E)** Illustration of the six time points at which TMS was delivered. The negative timings used to specify pre-movement TMS are referenced to each individual’s reaction time during the preceding block of trials. The times that elapsed between target onset and stimulation at each epoch therefore varied somewhat between subjects. CRT task: choice reaction time task; TMS: transcranial magnetic stimulation; FCR: flexor carpi radialis; ECRb: extensor carpi radialis brevis; ECU: extensor carpi ulnaris; FCU: flexor carpi ulnaris.

Each trial began with a circular warning sign displayed at the centre of the computer screen for between 1 s and 2 s before target appearance. Upon target appearance, participants were required to initiate cursor movement toward the target as quickly as possible. The targets appeared as a 10° wide wedge-shape that extended to 75 % of the distance from the origin to the edge of the computer screen (10 cm). The cursor gain was set such that 20 N was required to reach the edge of the screen. Participants were given post-trial feedback to encourage consistent movement times of 150 ms to 250 ms, with movement time defined as the time taken for the cursor to move from 10 % to 90 % of the target distance. A successful acquisition of target was cued with two high-pitched tones (500 ms, 800 Hz sinusoid) after the cursor remained within a 10% radius from the centre of target for 10 ms.

The four targets appeared in a random order within every cycle of four trials to prevent anticipation. Inter-trial intervals were 2 s. Participants completed 8 blocks of 72 trials (each block: 4 targets x 18 trials per target) of the choice reaction task during the entire experiment in a randomised order. In four blocks, TMS was delivered only when the target was presented at 0°, and for the other four blocks only upon presentation of the 180° target (i.e. 144 TMS trials per subject). TMS was delivered to the left motor cortex at target appearance, before the predicted movement onset time (−100 ms, −75 ms, −50 ms and −25 ms; relative to mean reaction time of the preceding block) and at movement onset (Figure 1C). Stimuli at movement onset were triggered by the onset of EMG activity in the relevant prime movers for a given target location.

### Experimental setup

Participants sat in front of a computer screen located approximately 1.2 m away at eye level (Figure 1D). The left and right forearms were secured into a custom-made manipulandum, described previously (de Rugy *et al*., 2012), which allowed passive rotation of the forearm between neutral (midway between pronation and supination) and pronation. Both elbows were kept at 110° with the forearm parallel to the table and supported by the manipulandum. The wrists were fixed by a series of twelve adjustable metal clamps contoured around the metacarpal-phalangeal joints and around the wrist proximal to the radial head. Wrist forces in abduction-adduction and flexion-extension directions were recorded via a six degree-of-freedom force transducer (JR3 45E15A-163-A400N60S, Woodland, CA) attached to each manipulandum. Force data were sampled at a rate of 2 kHz via two 16-bit National Instruments A/D boards (NI BNC2090A, NI USB6221, National Instruments Corporation, USA). The forces exerted in flexion-extension and abduction-adduction directions were displayed as a cursor that moved in two dimensions (for the neutral posture: x = flexion-extension, y = abduction-adduction) on the computer screen via a custom written Labview program (LabView2009, National Instrument, USA). The program also controlled the timing of TMS delivery.

### Transcranial magnetic stimulation

Single-pulse TMS was delivered via a 55 mm mid-diameter figure-of-eight magnetic coil (Magstim 200, Magstim, UK) over the forearm area of the left motor cortex. The magnetic coil was held tangentially on the scalp with the handle pointing backwards and 45° away from mid-sagittal axis to induce a posterior-anterior current direction in the brain. The resultant wrist forces induced by the TMS in abduction-adduction and flexion-extension directions were calculated and displayed on the computer screen. The optimal site eliciting the largest and most consistent TMS-evoked muscle twitches from the forearm was marked with a felt-tipped pen to ensure consistent stimulation throughout the experiment. The testing intensity was defined as an intensity that elicited a resultant force of magnitude between 0.5 to 1 N. The median stimulator intensity required was 55% of the maximal output, and ranged between 50 and 67% across subjects. To ensure evoked twitches were oriented in a near vertical orientation, we positioned the forearm either in a pronated or a neutral posture. We showed previously (Chye *et al*., 2013) that the direction of TMS-evoked twitches follows the muscles when the wrist is rotated between pronated and neutral positions. For example, if a twitch evoked from the right wrist in the neutral position is oriented horizontally rightwards (extension), then twitch direction will point vertically upwards when the wrist is rotated to a pronated position (still extension). It was therefore possible to ensure that each individual participant’s twitch directions at baseline were near vertical simply by repositioning their hand to a pronated or neutral position prior to the start of the experiment.

### Surface electromyography recordings

Electromyography (EMG) signals were recorded from flexor carpi radialis (FCR), flexor carpi ulnaris (FCU), extensor carpi radialis brevis (ECRb) and extensor carpi ulnaris (ECU) muscles of the both arms. Standard skin preparation was performed after the muscles were located and marked. Bipolar Ag/AgCl surface electrodes placed on the belly of the forearm muscles with an inter-electrode distance of 2 cm (centre to centre). The EMG signals were amplified with a gain of 500 ~ 1000 with Grass P511 amplifiers (Grass Instruments, AstroMed, West Warwick, RI) and band-pass filtered (10 Hz – 1 kHz).

### Data analysis

TMS-evoked twitch angles, TMS-evoked twitch magnitudes, motor evoked potential (MEP) amplitudes, and active limb premotor reaction times were extracted from force and EMG time-series data offline via a custom-written Matlab program (Mathworks, Natick, USA). Individual force and EMG traces were inspected visually, and trials containing voluntary EMG on muscles in the passive limb, or with force traces contaminated by postural or voluntary force were removed manually. For example, the onset of a twitch following an MEP onset occurs at latencies between 20 ms and 40 ms. Therefore, any force traces including substantial transients with onsets before or after this time range were removed from subsequent analysis. A total of 13 % of all TMS trials were excluded from analysis. Twitch and MEP responses obtained from the passive limb were grouped into bins of 40 ms width in relation to the timing of movement onset in the active limb, in order to assess the evolution of crossed effects during movement preparation and execution. Note that responses were grouped with respect to the actual movement time of the active limb on each trial, rather than the intended timing based on the average pre-motor reaction time from previous blocks. All data were presented as mean [95% confidence interval] unless stated otherwise. Statistical significance was set at the 0.05 level.

#### Twitch responses

Twitch angle and twitch magnitude at the time of peak resultant force were normalised by subtracting the mean values obtained from stimulation at the time of target appearance (referred to as baseline). The differences in twitch angles from baseline determined the reference frame according to which the twitches shifted, as illustrated in Figure 1A. Note that the baseline twitch direction was, by design, close to the vertical axis, and that TMS was delivered for horizontal targets only. Positive deviations of twitch angle were defined as when the directions of twitches evoked during movement preparation were closer to the target direction, defined in muscle space, than the baseline twitch direction. Thus, positive angles reflect that twitch direction in the passive limb “shifted” toward the pulling direction of the muscles homologous to those soon to be recruited in the active limb. Conversely, negative angle deviations were defined as when twitch direction shifted toward the movement direction defined in extrinsic space. Note also that a body-midline referenced coordinate system aligns with the muscle-based coordinate system (refer to Figure 1A).

#### MEP responses

The peak-to-peak MEP amplitudes were calculated for each muscle in both target conditions and normalised by subtracting the mean baseline MEP amplitude, and then by dividing by the peak MEP size recorded in that muscle (i.e. to either target direction at any stimulus timing). The MEP data for the four muscles were grouped across individuals according to whether each muscle was homologous to an agonist or antagonist for the movement performed with the active limb. Note that the specific muscles differed for participants who performed the task in pronated (n = 9) and neutral (n = 3) hand positions, since each muscle corresponded to different target directions in the two postures. For example, the ECR and FCR muscles were agonists, and the ECU and FCU were antagonists, when the left limb exerted force toward the 0° target (horizontal right target) when the forearm was pronated (refer to Figure 1E). Muscles were further categorised as “primary” or “secondary” on the basis of whether their pulling directions aligned with the baseline twitch directions (pulling directions were taken from de Rugy et al. 2012). Primary agonists were defined as muscles with pulling directions lying between the target direction and the baseline twitch direction. Secondary agonists were defined as muscles with pulling directions aligned with the target (i.e. within 90 degrees), but away from the baseline twitch direction (more than 90 degrees away). Antagonist muscles were similarly specified as primary or secondary. For example, with the forearms in a neutral posture and for a 0 degree target when the baseline twitch direction was near 90 degrees, the primary agonist was the FCR, the secondary agonist was the FCU, the primary antagonist was the ECR, and the secondary antagonist was the ECU.

#### Pre-motor reaction time

Pre-motor reaction time was defined as the interval between target appearance and the onset of the earliest EMG activity in the agonist muscles. The EMG activity threshold was defined as the amplitude that exceeded three times the standard deviation of the EMG signal prior to stimulus onset (Konrad, 2005). It has been reported that TMS of the active limb shortens reaction time when delivered during early motor preparation and delays reaction time when delivered during late motor preparation (Ziemann *et al*., 1997). To determine whether TMS delivered to the ipsilateral (“passive”) M1 alters reaction time, the (within subject) median premotor reaction times for trials in which TMS was delivered were compared with the non-stimulation trials for both target conditions. To assess the temporal evolution of any effects, the median pre-motor reaction times were also compared across six time points, based on the time from stimulus presentation to the (a-priori) estimated pre-motor reaction time: at target appearance, before movement onset (−100 ms, −75 ms, −50 ms and −25 ms) and at movement onset.

### Statistical analyses

All data were screened with the Shapiro-Wilk test for data normality. The twitch angle and pre-motor reaction time data were normally distributed (p > 0.05), and parametric analyses were performed. The Greenhouse-Geisser correction for degrees of freedom was applied when the assumption of sphericity was violated. Partial eta squared measures of effect size are reported for ANOVA effects, and Cohen’s d effect sizes were reported for pairwise contrasts. The twitch magnitude and MEP data were not normally distributed (p < 0.05), and thus a rank-transformation was performed prior to parametric analyses for these variables following Baguley (2012). Note that the statistical outcomes were almost identical if the analyses were performed on the un-transformed data. To determine whether there were significant changes from baseline for each time bin, independent Student’s t-tests against 0 were used for normally distributed variables, and sign tests were used for variables that were not normally distributed. Note that the p-values reported for tests against 0 are uncorrected for multiple comparisons, but all significant contrasts observed survive Bonferroni corrections. The same two-way repeated measures ANOVA model was applied to twitch magnitude data after rank transformation to determine the effect of time on twitch size.

## Results

### TMS evoked twitch angles for the resting right wrist

The direction of twitches evoked in the passive right wrist varied as a function of stimulation time relative to the onset of movement in the active left wrist, as illustrated by a main effect of TMS delivery time on the group mean change in twitch angle (F_(2.2,24.2)_ = 10.23, p = 0.0005, ηp^2^ = 0.48, Figure 2A), from a two-way repeated measures ANOVAs (target location [0 degree, 180 degree] x stimulus timing relative to movement onset [>120ms, 120-80ms, 80-40ms, 40-0ms, 0-40ms]). There was no main effect of target location (F_(1,11)_ = 0.01, p = 0.91, ηp^2^ = 0.001) or interaction between target location and TMS time (F_(2,21.8)_ = 0.52, p = 0.60, ηp^2^ = 0.04), suggesting that the time-varying effects were consistent irrespective of which direction of motion was performed by the right wrist. When pooled across the two targets, there was a significant shift from baseline in the twitch angle of approximately 28° [95% CI: 14.9, 40.8], toward the movement direction defined in body-referenced coordinates, for stimuli delivered after movement onset (time bin > 0 ms, t = 4.73, p = 0.0006, Cohen’s d = 1.64). Although there were no statistically significant changes in twitch angles during motor preparation, the apparent trend was also toward the movement direction defined in body-referenced coordinates from 80 ms prior to EMG onset (t value range: 1.49 to 1.68, p value range: 0.12 to 0.16, Cohen’s d range: 0.19 to 0.48).

**Figure 2.**
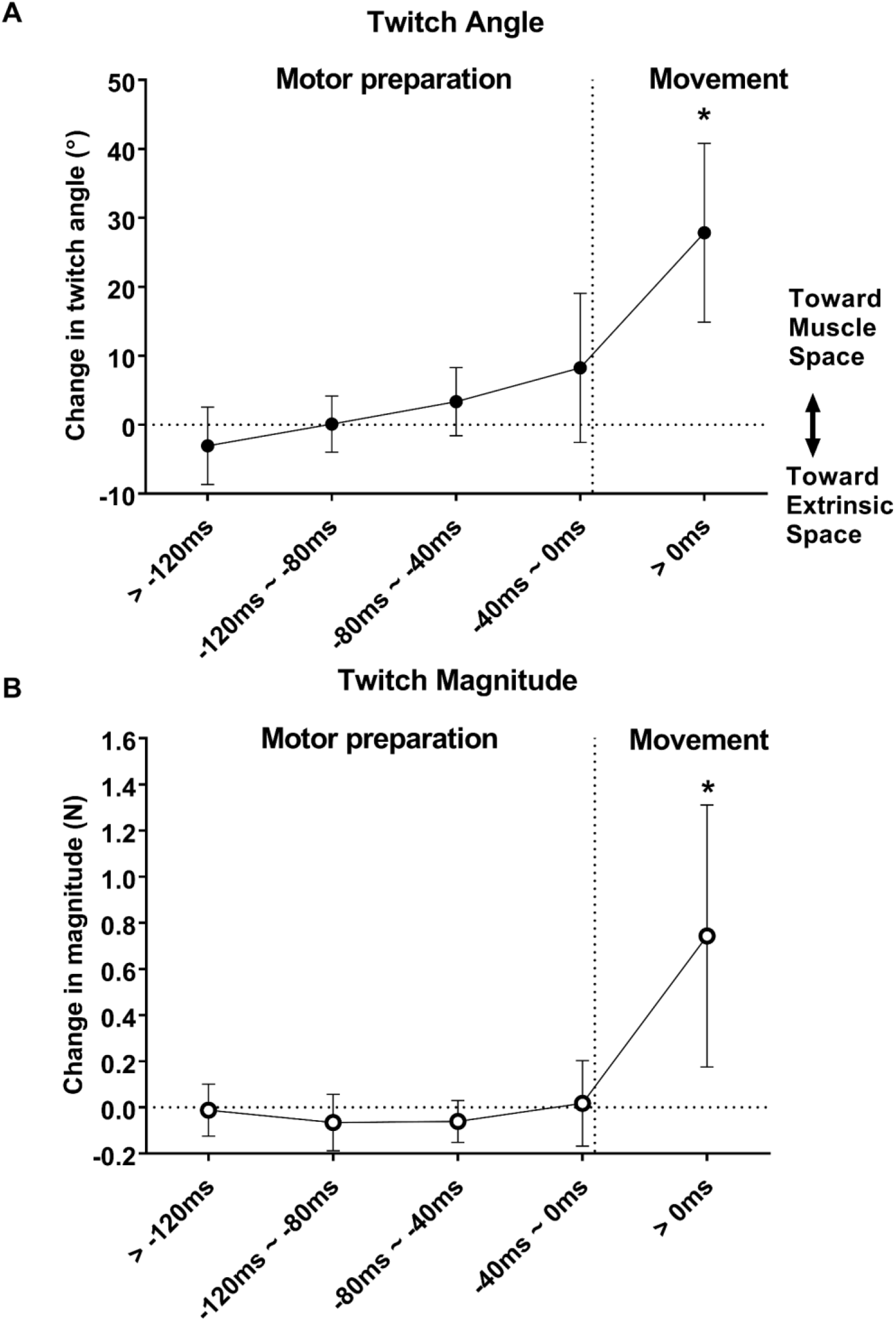
**(A)** Change in group mean (± 95% CI) twitch angles of the resting right wrist during motor preparation and post movement onset. Positive twitch angle changes depict that twitch direction shifted toward the opposite limb movement direction defined in muscle and body-midline space. Vertical dotted line depicts the movement onset of the active left wrist. Horizontal dotted line denotes the baseline. Symbol ‘*’ depicts significant difference from baseline (p < 0.05). **(B)** Change in group mean (± 95% CI) twitch magnitudes of the resting right wrist during motor preparation and post movement onset. The twitch magnitude increased during late motor preparation. Vertical dotted line depicts the movement onset of the active left wrist. Horizontal dotted line denotes the baseline. Symbol ‘*’ depicts significant difference from baseline (p < 0.05).

### TMS evoked twitch magnitudes for the resting right wrist

The time-course of changes in twitch magnitude appears grossly similar to that observed for twitch angles (Figure 2B). Thus, according to a two (0°, 180°) by five (stimulus timings) repeated measures ANOVA on the rank-transformed data, there was a significant main effect for stimulation time (F_(2.8,30.3)_ = 17.2, p = 2×10^-6^, ηp^2^ = 0.61), but no main effect for target location (F_(1,11)_ = 0.07, p = 0.79, ηp^2^ = 0.01) nor an interaction between target location and stimulation time (F_(2.7,29.3)_ = 0.83, p = 0.48, ηp^2^ = 0.07). When pooled across target locations, the twitch magnitude was significantly greater than baseline after movement onset (0.74 N change [95% CI: 0.2, 1.3], time bin > 0 ms, z = 2.59, p = 0.009, Cohen’s d = 1.41), and was not significantly different from baseline for any prior time-point (for all pairwise contrasts: z = 0.28, p = 0.77, Cohen’s d range: −0.02 to 0.03, Figure 2B). Note, that the effect sizes for twitch magnitude changes during preparation were negligible throughout preparation, in contrast to the increasing trend for twitch directions noted above.

### MEP amplitudes for the passive right wrist

The MEP amplitudes of all muscles in the passive limb in both target conditions were grouped and averaged according to whether their corresponding homologous muscles in the active limb were primary or secondary antagonists or agonists in the voluntary force pulse (see methods for a full explanation of muscle groupings). The absolute peak to peak amplitude of the MEPs obtained for the primary agonists in the control condition (i.e. when TMS was delivered at the time of target presentation) was 0.50 ± 0.15 mV (mean ± 95% CI). There were no significant main or interaction effects involving the factor primary versus secondary muscles, according to a rank transformed two (primary vs secondary) x two (agonist vs antagonist) x two (0 degree vs 180 degree target) x five (stimulation time) repeated measures ANOVA. This suggests that the time and agonist-antagonist muscle effects of motor preparation effects were similar irrespective of which muscles contributed most strongly to baseline twitches. Figure 3 shows the temporal evolution of changes in MEP size from baseline (n.b. any positive value represents an increase in MEP size from baseline), for the critical contrast of agonist versus antagonist muscle MEP size (pooled over primary and secondary muscles). There was a significant time x muscle (agonist vs antagonist) interaction (F_(2.1,22.6)_ = 74.6, p = 10^-6^, ηp^2^ = 0.87), indicating that MEPs increased to a greater extent in passive limb muscles that were homologous to the agonists rather than antagonists for the impending voluntary movement in the active limb. This implies that MEP changes in the passive limb were directionally tuned with respect to the upcoming movement according to body-referenced coordinates.

**Figure 3.**
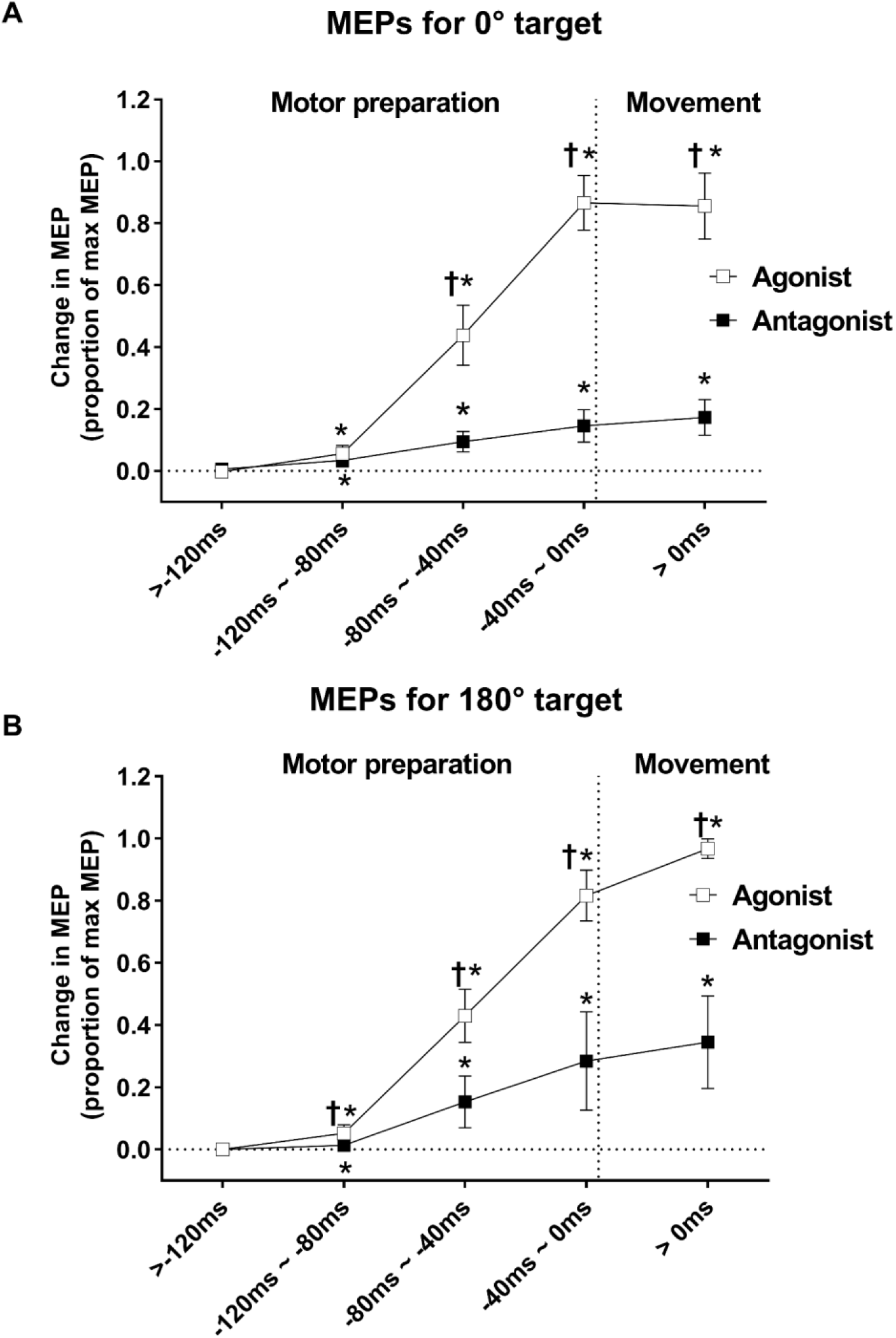
Changes from baseline in group mean (± 95% CI) MEP amplitudes for the antagonist and agonist muscles, during motor preparation and post movement onset. **(A)** Shows MEPs obtained when the active limb moved to the 0 degree target, **(B)** Shows MEPs recorded when the active limb moved to the 180 degree target. Vertical dotted line depicts the movement onset of the active left wrist. Horizontal dotted line denotes the baseline MEP size. MEP amplitude changes are expressed as a percentage of the maximal MEP size, observed for each participant and muscle, within a single time window and in any phase of the experiment. Black squares denote antagonist muscles. White squares denote agonist muscles. Symbol ‘*’ depicts significant difference from baseline (p < 0.05). Symbol ‘†’ depicts significant difference between the agonist and antagonist muscles (p < 0.05).

The changes in MEP size are shown in Figure 3, separately for movements to the 0 degree (towards the body midline) and 180 degree (away from the body midline) targets, since there was also a significant time x muscle x target interaction (F_(2.4,26)_ = 7.5, p = 0.002, ηp^2^ = 0.41). A Tukey post-hoc test on differences between changes in agonist and antagonist muscle MEP size showed that, for the 0 degree target, homologous agonist muscle MEP increases were greater than homologous antagonist muscle MEP increases at 80-40 and 40-0ms before EMG onset in the active limb, and after the movement onset (p = 0.0002 for all pairwise contrasts). For the 180 degree target, homologous agonist muscle MEP increases were greater than homologous antagonist muscle MEP increases at 120-80, 80-40, 40-0ms before EMG onset in the active limb, and after the movement onset (p = 0.0002 for all pairwise contrasts). Note that this pattern matches closely the qualitative pattern of muscle twitch direction changes over time, such that the greater increase in MEP size for agonists than for antagonists from ~100 ms before EMG onset reflects the general trend toward evoked twitches in intrinsic coordinates.

Figure 3 also shows that MEP amplitudes for muscles homologous to both antagonists and agonists were significantly larger than baseline (i.e. for stimuli delivered at the time of target presentation) from 120 ms before movement onset until 40 ms after movement onset (0 degree target: antagonist muscles: all z = 3.18, p = 0.001; agonist muscles: all z < 2.6, p < 0.01; 180 degree target: antagonist muscles: all z < 2.6, p < 0.01; agonist muscles: all z = 3.18, p = 0.001). However, MEP sizes in the passive limb were not different from baseline for stimuli delivered more than 120 ms before EMG onset in the active limb (0 degree target: antagonists: z = 0.29, p = 0.77, agonists: z = 0.29, p = 0.77; 180 degree target: antagonists: z = 0.29, p = 0.77, agonists: z = 0.87, p = 0.39).

### Pre-motor reaction time of the active left wrist

TMS of M1_contra_ is known to affect the reaction time of unilateral movements differently according to the time of its application. Here we found that the median pre-motor reaction times in the active left wrist were similar for stimulation and non-stimulation trials in both target directions, according to a two (stimulation vs no stimulation) x two (0 vs 180 degree targets) repeated measures ANOVA. (Figure 4A). Although the main effect for stimulation was marginal for statistical significance (F_(1,11)_ = 3.8, p = 0.07, ηp^2^ = 0.26), the nominal difference in premotor reaction time between stimulation and non-stimulation was small (i.e. reaction time for stimulation trials was ~ 6 ms shorter than non-stimulation trials). There was also no significant interaction between target direction and stimulation (F_(1,11)_ = 0.86, p = 0.37, ηp^2^ = 0.07). The median pre-motor reaction times for both target conditions were also similar irrespective of when TMS was delivered, i.e. from target appearance to voluntary force onset, according to a two (targets) x six (stimulation times) repeated measures ANOVA (Figure 4B). The timing of TMS with respect to the predicted time of opposite limb EMG had no significant effect on the reaction time for either target condition (main effect for stimulation timing: F_(5,55)_ = 1.77, p = 0.14, ηp^2^ = 0.14; interaction effect between stimulation timing and target direction: F_(5,55)_ = 1.54, p = 0.19, ηp^2^ = 0.12). Overall, the data suggest that TMS of M1_ipsi_ had little effect on the release time of motor commands to the active limb.

**Figure 4.**
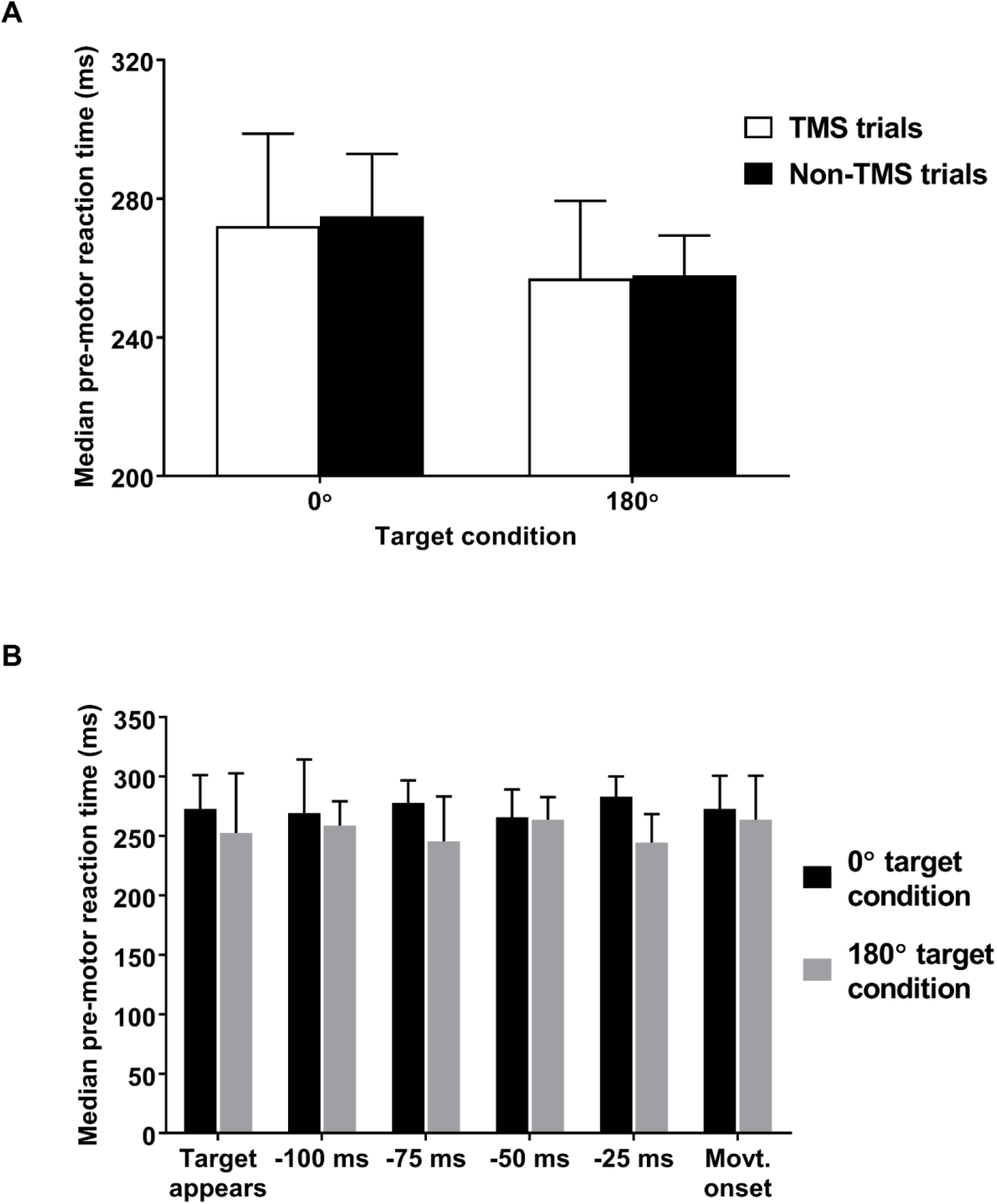
**(A)** Group means (± 95% CI) of median reaction times for each individual for the 0° and 180° target conditions, with (white bars) and without (black bars) TMS. **(B)** Group means (± 95% CI) of median reaction times for each individual for trials with TMS in each target condition at target appearance, at times prior to the estimated movement onset time (−100 ms, −75 ms, −50 ms and −25 ms) and at the estimated movement onset time. Black bars depict pre-motor reaction time for 0° target condition. Grey bars depict pre-motor reaction time for 180° target condition.

## Discussion

The current data show that the excitability of corticospinal projections to a passive limb is modulated in a directionally-specific manner during preparation for action with the opposite limb. Twitch directions, twitch magnitudes, and MEP amplitudes remained unchanged during early motor preparation (i.e. more than 120 ms prior to EMG onset), but directionally specific changes in MEP amplitudes were apparent from then onwards. There was a greater increase in the excitability of projections to muscles homologous to the agonists for the voluntary force pulse performed with the opposite limb than for muscles homologous to antagonists. Although there was a qualitatively similar trend for TMS-evoked twitches to shift towards the intrinsically defined target direction during late preparation, the effect was not statistically significant until the onset of voluntary EMG. Thus, the excitability of ipsilateral M1, and/or its downstream projections, reflect facilitation of neurons associated with the pulling direction of the muscles homologous to the prime movers in the active limb, and to the direction of the impending voluntary action relative to the body midline.

### Time-course of changes in TMS-evoked responses in the passive limb

Little modulation of corticospinal excitability was observed in the resting right wrist for TMS delivered between target appearance and 120 ms preceding movement onset. This is consistent with reports that corticospinal excitability changes little from baseline prior to 100 ms prior to movement, either in the responding hand (Chen *et al*., 1998; Leocani *et al*., 2000; Nikolova *et al*., 2006) or the resting hand (Leocani *et al*., 2000; Sommer *et al*., 2001; McMillan *et al*., 2006; van Elswijk *et al*., 2008). Here we found a general increase in MEP amplitudes for all of the muscles we sampled in the resting right wrist during late motor preparation. Such a late increase in responsiveness might be associated with reductions in interhemispheric inhibition or short-interval intracortical inhibition that have been reported during unilateral movement (Muellbacher *et al*., 2000; Soto *et al*., 2010; Howatson *et al*., 2011; Uehara *et al*., 2013; Reissig *et al*., 2014). MEP amplitude increases were greater for muscles homologous to the prime movers in the effector limb than for muscles corresponding to antagonists, however, suggesting that the earliest excitability changes in M1_ipsi_ are tuned to the impending pattern of muscle activation in the effector limb. Although caution is required when interpreting compound motor action potentials obtained via surface electrodes placed over muscles in the forearm, because the records are not selective to individual muscles (Selvanayagam *et al*., 2012), the fact that the pattern of excitability reversed for the two different targets (i.e. antagonists for movements to one target are antagonists for movements to the alternative target) strongly suggests a genuine directionally-specific effect.

Although MEP changes were observed from 120 ms before the onset of voluntary EMG in the active limb, the direction of twitches evoked in the passive limb did not shift significantly toward the direction of the impending contralateral force pulse until EMG onset. Although the general trend of twitch direction changes qualitatively matched the MEP changes, it appears that the small MEP differences observed between agonist and antagonist muscle responses were insufficient to generate reliable mechanical effects. It is also notable that directional changes in evoked twitches in the passive limb appeared later than previously reported for the active limb (e.g. 75-15ms, depending on the movement direction (Sommer *et al*., 2001; van Elswijk *et al*., 2008)). Nonetheless, it seems clear that substantial directionally-specific changes in TMS-evoked responses emerge in both limbs later than directional tuning in individual M1 neurons (Georgopoulos *et al*., 1989; Georgopoulos & Pellizer, 1995; Evarts, 2011), or task relevant modulations in M1 population dynamics (Ames *et al*., 2014); that is, more than 100 ms prior to movement onset. Kaufman et al. (2014) showed how preparatory cortical activity, such as directionally tuned activity in individual neurons, can occur without causing premature movement onset by reciprocal interactions that can be observed at the level of population dynamics. In this case, it would appear that the measure of premotor activity provided by TMS reflects corticospinal ensemble dynamics associated with “output-potent” rather than “output-null” states.

### Reference frame of directional shifts in TMS-evoked twitches

In order to move the limbs toward visual targets, the CNS must transform information about task goals from retinotopic to muscle-based coordinates (Andersen *et al*., 1993; Buneo & Andersen, 2006; Sabes, 2011). Previous work indicates that neural activity related to preparation and execution of wrist movements in M1_contra_ reflects both intrinsic, muscle-based movement parameters, and extrinsic directional movement parameters (N.B. the latter are indistinguishable from eye-based coordinates in many experimental designs; e.g. Kakei et al. (1999)). In contrast, the activity in M1_ipsi_ during unilateral movement appears to reflect joint– or muscle-based movement parameters to a greater degree than extrinsic spatial parameters (Ganguly *et al*., 2009). Although these studies provide compelling evidence that there is activity in both motor cortices that is related to impending movement, it is not clear whether such movement related activity is directly responsible for generating motor commands, as projections from the relevant neurons to the spinal cord were not verified (e.g. via spike-triggered averaging). Our current data suggest that excitability of the sub-population of neurons in M1_ipsi_ that is responsive to TMS, and that directly influences corticospinal output, reflects unilateral movement toward visuospatial targets according to muscle or body midline referenced coordinates. Such a representation could be the product of task-dependent excitability changes within M1_ipsi_, or at any point in the neuraxis between the M1 output cells and the motoneurons projecting to wrist muscles.

The correspondence of responses to stimulation of M1_ipsi_ with body-referenced movement parameters agrees with the general conclusions of Duque et al. (2005), who examined MEP responses to TMS of M1_ipsi_ in a context in which both bimanual and unimanual trials were randomly intermingled. In Duque’s et al. experiment, the required direction of movement was specified by symbolic cues, rather than the location of a visual target in extrinsic space, and TMS was applied from 100-40ms prior to EMG onset. However, these authors observed diminution of MEPs in an intrinsic hand muscle when its action mirrored the direction of motion of the opposite limb. In our study, however, there was a generalised increase of MEP amplitude within a similar time window. These changes had directionally-specific characteristics within the final 120 ms before movement onset. Our supposition is that the absence of diminution of MEP amplitude during any phase of movement preparation in the current study relates to the fact that the passive right hand was never required to move. This may have obviated the need for the suppression of latent responses that otherwise occurs in choice (of limb) reaction time tasks (see Bestmann and Duque (2015) for review). The late, muscle-specific facilitation that we observed corresponds to results obtained during execution of unilateral tonic (Hortobagyi *et al*., 2003; Perez & Cohen, 2008) and ballistic (Carroll *et al*., 2008; Carson & Ruddy, 2012) contractions, under conditions devoid of task uncertainty.

Our results differ from those of McMillan et al (2006), however, who found facilitation of MEP amplitudes from M1_ipsi_ that was consistent with the extrinsic line of action of the active limb. In that study, participants had to select which hand to use to make a movement toward a pre-specified target, based on a symbolic cue that also served as the imperative stimulus. It seems likely that this design would have prompted preparation for action with both limbs, and that the observed pattern of extrinsically-referenced MEP facilitation following stimulation of M1_ipsi_ might have reflected preparatory activity that was necessary to produce a rapid response in the alternative case in which the opposite limb would be specified.

In our design, a body-referenced representation of the impending movement with the ipsilateral limb is consistent with the notion that, prior to a strictly unilateral action, the excitability of projections from M1_ipsi_ represents the state of the active limb, rather than subliminal planning for the passive limb. This is because subliminal planning for the passive limb would necessarily be in extrinsic coordinates in our design, because a rightward movement would be needed to hit the rightward target with either limb. However, it is important to note that our data do not discriminate whether or not information about the state of the opposite (active) limb is represented in a muscle-based or a body-midline reference frame, or a combination of these. Further experiments are required to resolve this issue. A representation of the state of the active limb might serve a number of functional purposes. For example, information about the state of the ipsilateral limb would facilitate bimanual coordination in the event that the passive limb is required to produce an unexpected action during the target-directed movement. Representation of the current and future state of the active limb in both M1s might also make motor preparation robust to transient unilateral perturbations in cortical state, since both motor behavior and task-relevant neural population dynamics are rapidly restored after contralateral, but not bilateral, optogenetic silencing of premotor cortex in rodents (Li *et al*., 2016). Finally, activity in M1_ipsi_ might even play a role in actively shaping motor preparation in M1_contra_ via transcallosal interactions (Bianki & Makarova, 1980).

The increased responsiveness of M1_ipsi_ to cortical stimulation may relate to a subset of the mechanisms that can in some circumstances give rise to the expression of “mirror movements”. These are characterized by inadvertent contraction of muscles on one side of the body when the opposite limb produces an action that is by intention unilateral (see Carson, 2005 for review). It should be emphasised that in the present study, the participants produced relatively modest levels of contraction that did not cause fatigue. In this context, the state of some fraction of pathways projecting to the spinal cord, and then to muscles of the opposite limb was assessed using suprathreshold TMS i.e. at an intensity sufficient to elicit a descending corticospinal volley and a twitch response. There were no mirror movements per se. Although mirror movements are associated with a range of neurological conditions (Farmer *et al*., 1990; Galléa *et al*., 2011; Mayston et al. 1997), they can also occur in normally developing children (Abercrombie *et al*., 1964; Connolly & Stratton, 1968; Mayston *et al*., 1999; Wolff *et al*., 1983). In healthy adults, mirror movements are obtained most readily in the context of maximal effort movements or in performing contractions to the point of fatigue (Aranyi & Rosler, 2002; Hopf *et al*., 1974; Todor & Lazarus, 1986). In such cases, the inadvertent activity (registered either as movements or via electromyography) necessarily arises from bilateral interactions at multiple sites along the neuraxis (see Carson 2005 for review) i.e. extending beyond those sampled by the experimental techniques used in the present study. It cannot be assumed that the directional characteristics of the ipsilateral changes in neural excitability that occur across multiple networks in circumstances of maximal voluntary effort, all adhere to the principles elucidated in the current investigation. Perhaps most obviously, in such conditions the specificity with which muscles are recruited declines dramatically, and as a consequence joint torques and displacements in directions other than those intended are frequently observed (e.g. Carson & Riek, 2001; Kelso *et al*., 1993). This may account, at least in part, for a report that the inadvertent activity in muscles of the passive limb during isometric contractions sustained to the point of fatigue, gave rise to forces that were aligned to a greater degree with those applied by the active limb, than with the line of action of a principal muscle engaged in the task (Post *et al*., 2009). Further work is evidently required to determine the manner in which these differing constraints are integrated during actions performed in the course of daily living.

### Reaction time effects of M1_ipsi_ stimulation

Unilateral reaction time is modulated by TMS of the motor cortex contralateral to the active limb in a timing-dependent manner. Specifically, reaction time is shortened by approximately 50 to 75 ms when stimuli are delivered to the M1_contra_ at early stages of movement preparation (Leocani *et al*., 2000; McMillan *et al*., 2006; Nikolova *et al*., 2006; van Elswijk *et al*., 2008; Michelet *et al*., 2010; Soto *et al*., 2010), but lengthened by 50 – 80 ms when TMS is applied close to the expected movement onset (Ziemann *et al*., 1997; Leocani *et al*., 2000; McMillan *et al*., 2006; Nikolova *et al*., 2006; Michelet *et al*., 2010; Soto *et al*., 2010). In contrast, our current data indicate that premotor reaction time is similar during early and late motor preparation regardless of the timing of TMS applied to M1_ipsi_. This suggests that TMS of M1_ipsi_ has little effect on the time of release of motor commands during unilateral movement.

## Conclusion

The current results show that, during a unimanual choice reaction time task, directionally specific excitability changes within circuits projecting to the passive limb reflect the impending action in body referenced rather than extrinsic coordinates. This is consistent with the interpretation that ipsilateral motor cortical activity prior to unilateral action reflects the current or future state of the opposite limb, rather than subliminal planning for the passive limb.

## References

Abercrombie ML, Lindon RL & Tyson MC. (1964). Associated Movements in Normal and Physically Handicapped Children. Dev Med Child Neurol 6, 573–580.

Ames CK, Ryu SI & Shenoy KV. (2014). Neural dynamics of reaching following incorrect or absent motor preparation. Neuron 81, 438–451.

Andersen RA, Snyder LH, Li C-S & Stricanne B. (1993). Coordinate transformations in the representation of spatial information. Curr Opin Neurobiol 3, 171–176.

Aranyi Z & Rosler KM. (2002). Effort-induced mirror movements. A study of transcallosal inhibition in humans. Exp Brain Res 145, 76–82.

Baguley T. (2012). Serious stats: A guide to advanced statistics for the behavioral sciences. Palgrave Macmillan.

Bestmann S & Duque J. (2015). Transcranial magnetic stimulation: Decomposing the processes underlying action preparation. Neuroscientist, 1–14.

Bianki VL & Makarova IA. (1980). Transcallosal modulation of a focus of maximal activity in the motor cortex. Neurosci Behav Physiol 10, 76–82.

Buneo CA & Andersen RA. (2006). The posterior parietal cortex: sensorimotor interface for the planning and online control of visually guided movements. Neuropsychologia 44, 2594–2606.

Bütefisch C, Revill K, Shuster L, Hines B & Parsons M. (2014). Motor demand-dependent activation of ipsilateral motor cortex. J Neurophysiol 112, 999–1009.

Carroll TJ, Lee M, Hsu M & Sayde J. (2008). Unilateral practice of a ballistic movement causes bilateral increases in performance and corticospinal excitability. J Appl Physiol 104, 1656–1664.

Carson RG. (2005). Neural pathways mediating bilateral interactions between the upper limbs. Brain Res Brain Res Rev 49, 641–662.

Carson RG & Riek S. (2001). Changes in muscle recruitment patterns during skill acquisition. Experimental Brain Research 138, 71–87.

Carson RG & Ruddy K. (2012). Vision modulates corticospinal suppression in a functonally specific manner during movement of the opposite limb. J Neurosci 32, 646–652.

Chen R, Yaseen Z, Cohen LG & Hallett M. (1998). Time course of corticospinal excitability in reaction time and self-paced movements. Ann Neurol 44, 317–325.

Chiou SY, Wang RY, Roberts R, Wu YT, Lu CF & Liao KK. (2014). Fractional anisotropy in corpus callosum is associated with facilitation of motor representation during ipsilateral hand movements. Plos One 9, 1–9.

Chye L, Riek S, de Rugy A & Carroll T. (2013). Do TMS-evoked twitches shift towards a ballistic training direction in extrinsic or muscle-based coordinates? In Neuroscience Conference. San Diego, California.

Connolly K & Stratton P. (1968). Developmental changes in associated movements. Dev Med Child Neurol 10, 49–56.

de Rugy A, Davoodi R & Carroll TJ. (2012). Changes in wrist muscle activity with forearm posture: implications for the study of sensorimotor transformations. J Neurophysiol 108, 2884–2895.

Duque J & Ivry RB. (2009). Role of corticospinal suppression during motor preparation. Cereb Cortex 19, 2013–2024.

Duque J, Lew D, Mazzocchio R, Olivier E & Ivry RB. (2010). Evidence for two concurrent inhibitory mechanisms during response preparation. J Neurosci 30, 3793–3802.

Duque J, Mazzocchio R, Dambrosia J, Murase N, Olivier E & Cohen LG. (2005). Kinematically specific interhemispheric inhibition operating in the process of generation of a voluntary movement. Cereb Cortex 15, 588–593.

Duque J, Mazzocchio R, Stefan K, Hummel F, Olivier E & Cohen LG. (2008). Memory formation in the motor cortex ipsilateral to a training hand. Cereb Cortex 18, 1395–1406.

Evarts EV. (2011). Role of motor cortex in voluntary movements in primates. Compr Physiol, 1083–1120.

Farmer SF, Ingram DA & Stephens JA. (1990). Mirror movements studied in a patient with Klippel-Feil syndrome. J Physiol 428, 467–484.

Gallea C, Popa T, Billot S, Meneret A, Depienne C & Roze E. (2011). Congenital mirror movements: a clue to understanding bimanual motor control. J Neurol 258, 1911–1919.

Ganguly K & Poo M. (2013). Activity-dependent neural plasticity from bench to bedside. Neuron 80, 729–741.

Ganguly K, Secundo L, Ranade G, Orsborn A, Chang E, Dimitriov D, Wallis JD, Barbaro NM, Knight RT & Carmena JM. (2009). Cortical representation of ipsilateral arm movements in monkey and man. J Neurosci 29, 12948–12956.

Georgopoulos AP, Lurito JT, Petrides M, Schwartz AB & Massey JT. (1989). Mental rotation of the neuronal population vector. Science 243, 234–236.

Georgopoulos AP & Pellizer G. (1995). The mental and the neural: psychological and neural studies of mental rotation and memory scanning. Neuropsychologia 33, 1531–1547.

Griffin DM, Hoffman DS & Strick PL. (2015). Corticomotoneuronal cells are “functionally tuned”. Science 350, 667–670.

Groppa S, Oliviero A, Eisen A, Quartarone A, Cohen LG, Mall V, Kaelin-Lang A, Mima T, Rossi S, Thickbroom GW, Rossini PM, Ziemann U, Valls-Solé J & Siebner HR. (2012). A practical guide to diagnostic transcranial magnetic stimulation: report of an IFCN committee. Clin Neurophysiol 123, 858–882.

Hinder MR, Schmidt MW, Garry MI & Summers JJ. (2010). The effect of ballistic thumb contractions on the excitability of the ipsilateral motor cortex. Exp Brain Res 201, 229–238.

Hopf HC, Schlegel HJ & Lowitzsch K. (1974). Irradiation of voluntary activity to the contralateral side in movements of normal subjects and patients with central motor disturbances. Eur Neurol 12, 142–147.

Hortobagyi T, Taylor JL, Petersen NT, Russell G & Gandevia SC. (2003). Changes in segmental and motor cortical output with contralateral muscle contractions and altered sensory inputs in humans. J Neurophysiol 90, 2451–2459.

Howatson G, Taylor MB, Rider P, Motawar BR, McNally MP, Solnik S, DeVita P & Hortobagyi T. (2011). Ipsilateral motor cortical responses to TMS during lengthening and shortening of the contralateral wrist flexors. Eur J Neurosci 33, 978–990.

Kakei S, Hoffman DS & Strick PL. (1999). Muscle and movement representations in the primary motor cortex. Science 285, 2136–2139.

Kalaska JF & Crammond DJ. (1992). Cerebral cortical mechanisms of reaching movements. Science 255, 1517–1523.

Kaufman MT, Churchland MM, Ryu SI & Shenoy KV. (2014). Cortical activity in the null space: permitting preparation without movement. Nat Neurosci 17, 440–448.

Kelso JAS, Buchanan JJ, Deguzman GC & Ding M. (1993). Spontaneous Recruitment and Annihilation of Degrees of Freedom in Biological Coordination. Phys Lett A 179, 364–371.

Kertesz A & Geschwind N. (1971). Patterns of pyramidal descussation and their relationship to handedess. Arch Neurol 24, 326–332.

Kim SG, Ashe J, Hendrich K, Ellermann J, Merkle H, Ugurbil K & Georgopoulos AP. (1993). Functional magnetic resonance imaging of motor cortex: Hemispheric asymmetry and handedness. Science 261, 615–617.

Konrad P. (2005). The ABC of EMG: A practical introduction of kinesiological electromyography. Noraxon INC, USA.

Lee M, Hinder MR, Gandevia SC & Carroll TJ. (2010). The ipsilateral motor cortex contributes to cross-limb transfer of performance gains after ballistic motor practice. J Physiol (Lond) 588, 201–212.

Leocani L, Cohen LG, Wassermann EM, Ikoma K & Hallett M. (2000). Human corticospinal excitability evaluated with transcranial magnetic stimulation during different reaction time paradigms. Brain 123, 1161–1173.

Levy J. (2013). Cerebal asymmetry and the psychology in man. In The brain and psychology, ed. Wittrock MC. Academic Press.

Li N, Daie K, Svoboda K & Druckmann S. (2016). Robust neuronal dynamics in premotor cortex during motor planning. Nature 532, 459–464.

McMillan S, Ivry RB & Byblow WD. (2006). Corticomotor excitability during a choice-hand reaction time task. Exp Brain Res 172, 230–245.

Mayston MJ, Harrison LM, Quinton R, Stephens JA, Krams M & Bouloux PM.(1997). Mirror movements in X-linked Kallmann’s syndrome. I. A neurophysiological study. Brain 120 (Pt 7), 1199–1216.

Mayston MJ, Harrison LM & Stephens JA. (1999). A neurophysiological study of mirror movements in adults and children. Ann Neurol 45, 583–594.

Michelet T, Duncan GH & Cisek P. (2010). Response competition in the primary motor cortex: corticospinal excitability reflects response replacement during simple decisions. J Neurophysiol 104, 119–127.

Muellbacher W, Facchini S, Boroojerdi B & Hallett M. (2000). Changes in motor cortex excitability during ipsilateral hand muscle activation in humans. Clin Neurophysiol 111, 344–349.

Nikolova M, Pondev N, Christova L, Wolf W & Kossev AR. (2006). Motor cortex excitability changes preceding voluntary muscle activity in simple reaction time task. Eur J Appl Physiol 98, 212–219.

Oldfield RC. (1971). Assessment and analysis of handedness – Edinburgh Inventory. Neuropsychologia 9, 97-&.

Perez MA, Butler JE & Taylor JL. (2014). Modulation of transcallosal inhibition by bilateral activation of agonist and antagonist proximal arm muscles. J Neurophysiol 111, 405–414.

Perez MA & Cohen LG. (2008). Mechanisms underlying functional changes in the primary motor cortex ipsilateral to an active hand. J Neurosci 28, 5631–5640.

Post M, Bakels R & Zijdewind I. (2009). Inadvertent contralateral activity during a sustained unilateral contraction reflects the direction of target movement. J Neurosci 29, 6353–6357.

Reissig P, Garry MI, Summers JJ & Hinder MR. (2014). Visual feedback-related changes in ipsilateral cortical excitability during unimanual movement: Implications for mirror therapy. Neuropsychological Rehabilitation, 1-22.

Rossi S, Hallett M, Rossini PM & Pascual-Leone A. (2011). Screening questionnaire before TMS: an update. Clin Neurophysiol 122, 1686.

Sabes PN. (2011). Sensory integration for reaching: models of optimality in the context of behavior and the underlying neural circuits. Prog Brain Res 191, 195–209.

Scott SH, Gribble P, Graham KM & Cabel DW. (2001). Dissociation between hand motion and population vectors from neural activity in motor cortex. Nature 413, 161–165.

Selvanayagam VS, Riek S & Carroll TJ. (2012). A systematic method to quantify the presence of cross-talk in stimulus-evoked EMG responses: Implications for TMS studies. J Appl Physiol 112, 259–265.

Siegel A & Sapru HN. (2011). Essential neuroscience. Wolters Kluwer Health/Lippincott Williams & Wilkins, Philadelphia.

Sommer M, Classen J, Cohen LG & Hallett M. (2001). Time course of determination of movement direction in the reaction time task in humans. J Neurophysiol 86, 1195–1201.

Soto O, Valls-Sole J & Kumru H. (2010). Paired-pulse transcranial magnetic stimulation during preparation for simple and choice reaction time tasks. J Neurophysiol 104, 1392–1400.

Todor JI & Lazarus JA. (1986). Exertion level and the intensity of associated movements. Dev Med Child Neurol 28, 205–212.

Uehara K, Morishita T, Kubota S & Funase K. (2013). Neural mechanisms underlying the changes in ipsilateral primary motor cortex excitability during unilateral rhythmic muscle contraction. Behav Brain Res 240, 33–45.

van Elswijk G, Schot WD, Stegeman DF & Overeem S. (2008). Changes in corticospinal excitability and the direction of evoked movements during motor preparation: a TMS study. BMC Neurosci 9.

Verstynen T & Ivry RB. (2011). Network dynamics mediating ipsilateral motor cortex activity during unimanual actions. J Cogn Neurosci 23, 2468–2480.

Wolff PH, Gunnoe CE & Cohen C. (1983). Associated Movements as a Measure of Developmental Age. Developmental Medicine and Child Neurology 25, 417–429.

Ziemann U, Tergau F, Netz J & Homberg V. (1997). Delay in simple reaction time after focal transcranial magnetic stimulation of the human brain occurs at the final motor output stage. Brain Res 744, 32–40.

